# Cell invasive amyloid assemblies from SARS-CoV-2 peptides can form multiple polymorphs with varying neurotoxicity

**DOI:** 10.1101/2024.05.16.594465

**Authors:** Oana Sanislav, Rina Tetaj, Metali, Julian Ratcliffe, William Phillips, Annaleise Klein, Ashish Sethi, Jiangtao Zhou, Raffaele Mezzenga, Sina Saxer, Mirren Charnley, Sarah Annesley, Nicholas P Reynolds

**Author notes:** **Conflict of Interest Statement:** The authors declare no conflict of interest. **Data Availability:** Upon acceptance of publication all data from this paper will be uploaded to a publicly accessible repository such as Zenodo. **Funding Statement:** This work was funded in part by the following: - The CASS foundation (#10053, ‘Determining the role of protein aggregation in COVID-19’) - The Australian Nuclear Science and Technology Organisation (ANSTO) (M19465) (M17173).

## Abstract

The neurological symptoms of COVID-19, such as memory loss, cognitive and sensory disruption (neuro-COVID) are well reported. These neurological symptoms frequently persist for months (post-acute sequalae of COVID-19 or PASC). The molecular origins of neuro-COVID and how it contributes to PASC are unknown, however a growing body of research highlights that the self-assembly of protein fragments from SARS-CoV-2 into amyloid nanofibrils may play a causative role. Previously, we identified two fragments from the proteins Open Reading Frame 6 (ORF6) and ORF10 that self-assemble into neurotoxic amyloid assemblies. Here we further our understanding of the self-assembly mechanisms and nano-architectures formed by these fragments as well as performing a more in-depth study of the biological responses of co-cultured neurons. By solubilising the peptides in a fluorinated solvent we eliminate insoluble aggregates in the starting materials (seeds) that change the polymorphic landscape of the assemblies. The resultant assemblies are dominated by structures with higher free energies (e.g. ribbons and amorphous aggregates) that are less toxic to cultured neurons. We also show the first direct evidence of cellular uptake by viral amyloids. This work highlights the importance of understanding the polymorphic behaviour of amyloids particularly in the context of neuro-COVID and PASC.

**Graphical Abstract for ToC:** The neurological symptoms of COVID-19 are likely to be, in part, caused by the aggregation of viral proteins into neurotoxic amyloid nanofibrils. Changes in aggregation conditions alters the balance of fibril structures formed (polymorphism), influencing their toxicity to a neuronal cell line. These findings increase our understanding of viral amyloids and highlight the importance of careful choice of experimental protocol when studying these systems.

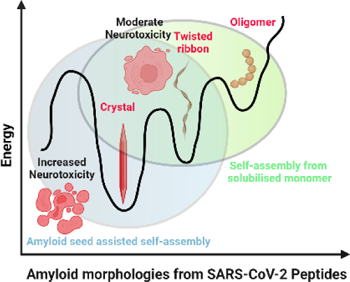

## 1. Introduction

Infection with SARS-CoV-2 and the associated disease (COVID-19) affects almost every organ in the body. The neurological symptoms of COVID-19 (sometimes referred to as neuro-COVID) are becoming more apparent.[1] Neuro-COVID symptoms include memory loss, cognitive issues, severe headaches, inflammation, anosmia and phantosmia (loss of smell and taste) and in rare cases haemorrhagic stroke.[2–5] These neurological symptoms are commonly reported in patients suffering from post-acute sequelae (PAS) of COVID-19 (PASC or long-COVID).[1] Due to their similarities to symptoms of neurodegenerative diseases (ND’s) such as Alzheimer’s (AD) and Parkinson’s (PD) [6, 7] a number of researchers around the world have begun to speculate that these symptoms might be caused by the build-up of toxic β-sheet rich amyloid assemblies in the central nervous system of infected patients.[8–14]

If amyloids are formed in the brain of neuro-COVID patients this begs the question, what is the source of these amyloids? Recent works have pointed to a number of potential viral amyloid protein culprits, including Spike[11], NSP11,[13] NCAP,[12] and in work previously published from our labs ORF6.[9] ORF6 as an amyloidogenic protein is a logical choice as a toxic culprit for the following reasons: ORF6 amyloids are known to be neurotoxic;[9] it is likely that monomeric ORF6 has an unstructured architecture (making aggregation of amyloidogenic fragments more feasible); ORF6 has been shown to possess increased cytotoxicity compared to other proteins expressed by SARS-CoV-2 infected cells;[15] it can block nucleocytoplasmic transport[16] and possesses significant immune silencing properties.[17, 18] All together ORF6 is a highly pathological protein and some of these pathologies may possess an amyloid aetiology.

It is important to understand in more detail how these viral protein assemblies can alter cellular responses and promote pathology. This knowledge can then be compared to previous studies on amyloids connected to ND’s, leading to a better understanding of disease progression in PASC and other infections due to viral amyloids. Previous works have shown that Amyloid-β (Aβ) aggregates are pathological to neuronal cells triggering cell death by a number of different mechanisms including, DNA damage, inhibition of ion-motive ATPases, and loss of Ca^+^ homoeostasis.[19] Further Aβ has been shown to cause impaired mitochondrial respiration in neuronal cells,[20, 21] and AD patients have significantly reduced mitochondrial respiration[22]. Conversely, work from our labs has shown that α-synuclein fibrils result in hyperactive mitochondrial respiration in the same cell lines (SH-SY5Y)[23]. The above works show that the effect amyloids have on mitochondrial respiration is complex and specific to individual proteins/polymorphs, therefore it is important to study in the context of viral amyloids.

In our previous work through a combination of bioinformatic and nanoanalytical studies we identified peptide fragments from the viral proteins ORF6 (ILLIIM) and ORF10 (RNYIAQVD) that assembled into a polymorphic array of amyloid architectures. ILLIIM almost exclusively formed crystalline structures with a well-defined cross-β structure,[9] and RNYIAQVD was formed a mixture of amyloid crystals and twisted ribbons. It is well established that amyloid crystals possess lower free energies (likely representing a global energy minima) than twisted and helical ribbons,[24, 25] (Figure 1) therefore we hypothesised that the added stability of the ILLIIM assemblies is connected to their increased cytotoxicity (slower cellular clearance of more stable amyloids leading to increased toxicity). In addition to polymorphism leading to different architectures and free energy levels, the self-seeding of pre-formed or undissolved amyloid aggregates is known to accelerate aggregation[26–28] likely resulting in a greater proportion of more stable assemblies possessing lower free energies.[9, 24, 25]To fully understand the relationship between amyloid free energy and cell pathology it is important to initiate self-assembly from dissolved samples free from seeds that may alter the resultant energy levels of the final assembly products. [9, 25, 29, 30]

**Figure 1:**
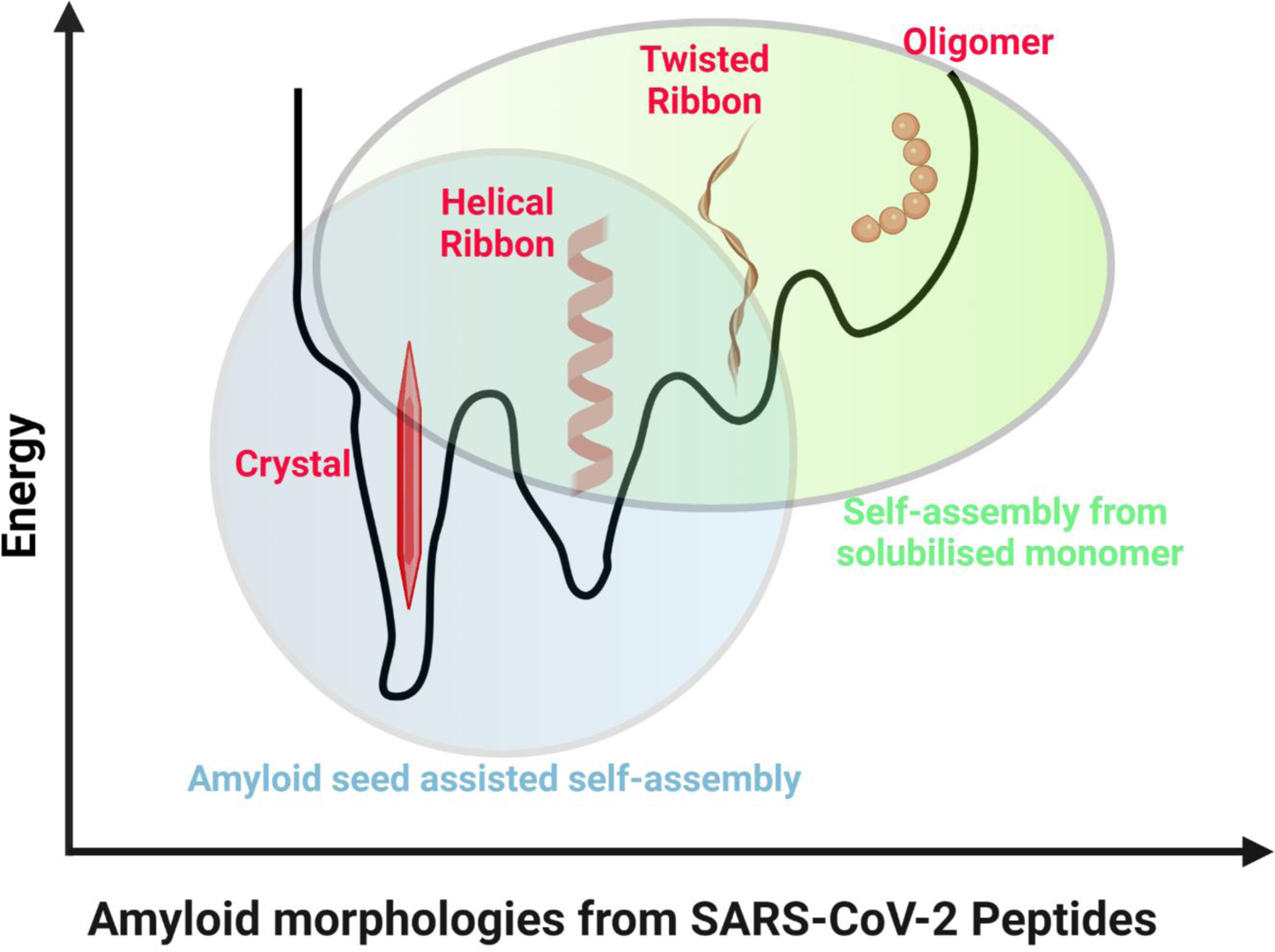
Amyloid assemblies formed from SARS-CoV-2 peptides are highly polymorphic, and the presence of pre-formed aggregates in the starting material (seeds) can promote the formation of lower energy assemblies (‘amyloid seed assisted self-assembly’) compared to solutions prepared in the absence of seeds (‘self-assembly from solubilised monomer).

To ensure the solubilisation of our two peptides we used the fluorinated solvent 1,1,1,3,3,3-hexafluoro-2-propanol (HFIP). HFIP is known to effectively disrupt hydrogen and other non-covalent bonds and prevent or reverse amyloid assembly.[11, 31] Using HFIP we were able to monitor the polymorphism of the two assembling peptides after buffer exchange and dilution into PBS. We showed that the resultant assembles possessed architectures consistent with amyloid assemblies with higher free energies[24, 32] than seen in Charnley et. al.[9] We observed that the amyloid assemblies are readily ingested by a neuronal cell line, and this caused significant cellular pathologies to cultured neurons and similarly to Charnley *et. al.*[9] were cytotoxic, although the extent of cytotoxicity was reduced due to the less stable assemblies formed.[9] In addition to shedding more light on the potential pathogenic roles of amyloid assemblies in neuro-COVID, this work highlights the importance of accurate control over the polymorphic assembly of amyloid proteins when inferring potential links to disease. Further it adds support to the hypothesis that amyloid seeding may promote additional pathologies in COVID-19, PASC and potentially other viral PAS.[10, 11]

## 2. Materials and Methods

### 2.1 Peptide Self-Assembly

NH_2_-ILLIIM-CO_2_H, Ac-RNYIAQVD-NH_2_ and NH_2_-AAAAAA-CO_2_H as a non-amyloid forming control (>98% pure) were purchased from GL Biochem Ltd (Shanghai, China). Ideally, it would have been preferred to have both viral peptides capped (N-terminus: Acetyl and C-terminus: Amide), as they would better represent small fragments of a larger peptide sequence. Due to the fact ILLIIM contains no charged sidechains, synthesising capped sequences to high purity would have been very challenging, therefore only the RNYIAQVD sequence remained capped and the ILLIIM sequence had regular carboxyl and amino termini. Both peptides were dissolved in 1,1,1,3,3,3-hexafluoro-2-propanol (HFIP, Sigma Aldrich) to stock solutions of 50 mg mL^-1^ and briefly vortexed. Stock solutions were diluted into PBS to their desired concentrations (0.2-1.5 mg mL^-1^), vortexed and allowed to self-assemble for 24 hours at room temperature in a laminar flow hood. During the first 3 hours of self-assembly the lid of the Eppendorf was left open to allow for the evaporation of the highly volatile HFIP. If this step was skipped, residual HFIP in the solution resulted in additional cytotoxicity of the samples. After assembly samples in PBS were kept sealed at 4 °C.

### 2.2 Fourier Transform-Infrared Microscopy (FT-IR)

Transmission FT-IR was performed at room temperature on the Infrared Microspectroscopy beamline at the Australian Synchrotron (AS) using the synchrotron radiation source and a Hyperion 3000 microscope with a liquid nitrogen-cooled Mercury Cadmium Telluride detector. Peptide assemblies in PBS at 1 mg mL^-1^ were prepared by depositing 2.5 µL of peptide solution onto a 13 mm × 0.5 mm BaF_2_ slide (Crystran Ltd). Samples were placed onto a hot plate at 37 °C and allowed to air dry. Single-point spectra were collected from visible deposits on the slide. Spectra were acquired using 64 co-added scans for both background and sample measurements at a resolution of 4 cm^-1^ employing a scanner velocity of 40 kHz, with a Blackman-Harris 3-term apodization function and zero filling factor of two. 40-point measurements were taken per sample using a beam size of 5.6 µm. Spectra were corrected for water vapour and CO_2_ using OPUS 8.0 (Bruker). Selected point spectra were averaged using OPUS 8 and the components of the amide I band of the each average spectrum were fitted using the non-linear curve fitting tool in OriginPro 2021.

### 2.3 Circular Dichroism Spectroscopy (CD)

CD spectroscopy was performed using an AVIV 410-SF CD spectrometer. Spectra were collected between 190 and 260 nm in PBS at 1 mg mL^-1^ using 1 mm quartz cuvettes with a step size of 0.5 nm and 2 s averaging time. Data were analysed using the BeStSel (Beta Structure Selection) method of secondary structure determination.[33]

### 2.4 Small and Wide Angle X-ray Scattering (SAXS/WAXS)

SAXS/WAXS experiments were performed at room temperature on the SAXS/WAXS beamline at the Australian Synchrotron (AS). Peptide assemblies in PBS prepared at 1 mg mL^−1^ were loaded into a 96-well PCR plate held on a robotically controlled x-y stage and transferred to the beamline via a quartz capillary connected to a syringe pump. The experiments used a beam wavelength of λ = 1.03320 Å^−1^ (12.0 keV) with dimensions of 300 µm × 200 µm and a typical flux of 1.2 × 10^13^ photons per second. 2D diffraction images were collected on a Pilatus 1M detector. SAXS experiments were performed at q ranges between 0.002 and 0.25 Å^−1^ and WAXS experiments were performed at a q range between 0.1 and 2 Å^−1^. These overlapping spectra provide a total q range of 0.002–2.2 Å^−1^. Spectra were recorded under flow (0.15 mL min^−1^) to prevent X-ray damage from the beam. Multiples of approximately 15 spectra were recorded for each time point (exposure time = 1 s) and averaged spectra are shown after background subtraction against PBS in the same capillary. Averaging of the spectra and background subtraction was performed using the software developed at the AS (ScatterBrain). Fitting of the SAXS spectra to form factors for flattened bicelle (ILLIIM) and flexible cylinder (RNYIAQVD) models was performed in SASView using average pre-determined fibril diameters calculated from AFM and TEM data.

### 2.5 Thioflavin T (ThT) Fluorescence Assay

Small volumes (∼0.5-2 μl depending on the final concentration desired) of stock solution of peptide in HFIP [50 mgmL^-1^] was transferred to an empty well in a glass bottomed black plastic 96-well plate. Using the microinjection device on the ClarioStar UV/Vis fluorometer a solution of ThT in PBS was injected into the same well making a final ThT concentration of 25 μM and a peptide concentration of between 0.05-2 mg mL^-1^. Immediately after injection the well plate was shaken for 30 s (double orbital shaking), and the first fluorescence measurement (*t* = 0 (excitation wavelength: 440 nm, emission wavelength: 482 nm) was recorded immediately after shaking, subsequent measurements were taken every 10 s until a plateau was reached.

### 2.6 Transmission Electron Microscopy (TEM)

Copper TEM grids with a formvar-carbon support film (GSCU300CC-50, ProSciTech, Qld, Australia) were glow discharged for 60 s in an Emitech k950x with k350 attachment. Then, 5 µL drops of sample suspension were pipetted onto each grid, allowed to adsorb for at least 30 s and blotted with filter paper. Two drops of 2% uranyl acetate were used to negatively stain the particles with excess negative stain removed by blotting with filter paper after 10 s each. Grids were then allowed to dry before imaging. Grids were imaged using a Joel JEM-2100 (JEOL (Australasia) Pty Ltd) transmission electron microscope equipped with a Gatan Orius SC 200 CCD camera (Scitek Australia).

### 2.7 Atomic Force Microscopy (AFM)

AFM imaging was performed on a Bruker Multimode 8 AFM and a Nanoscope V controller. Tapping mode imaging was used throughout, with antimony (n)-doped silicon cantilevers having approximate resonant frequencies of 150 kHz and spring constants of 5 Nm^−1^ (RTESPA-150, Bruker, U.S.A.). Freshly cleaved mica was first functionalized by 10 µL (3-Aminopropyl)triethoxysilane (1%) for 2 min, followed by rinsing with MQ water and drying under a nitrogen streamy. Then 10 µL aliquots of the peptide solution were drop cast onto functionalized mica disks and incubated for 2 min before gently rinsing in MQ water and drying under a nitrogen stream. All images were flattened using the nanoscope analysis software (Bruker, U.S.A.) and statistical analysis of the AFM images was performed using the open-source software FiberApp[34].

### 2.8 Cell Culture

Human-derived neuroblastoma cells (SH-SY5Y, ATCC Product Number: CRL-2266) were cultured in DMEM-F12 (Invitrogen) medium supplemented with 10 % (v/v) foetal bovine serum (FBS), 100 UmL^−1^ penicillin and 100 µgmL^−1^ streptomycin (Invitrogen, Carlsbad, CA). Cells were cultured at 37 °C in a humidified atmosphere containing 5 % CO_2_.

### 2.9 Cell Viability Assays

Cells were seeded into 96-well plates at 5 × 10^5^ cells per mL and incubated for 24 h to ensure good attachment to the surface. A stock solution of peptide assemblies (10 mg mL^−1^) was serially diluted to a concentration range found to be span a window of low-to-high toxicity in Charnley *et. al.*^9^ (160-20 μg mL^−1^) into DMEM-F12 media and seeded onto the SH-SY5Y cells and incubated for 48 h, cell viability was determined using 3-(4,5-dimethylthiazol-2-yl)2,5-diphenyltetrazolium bromide (MTT) (Sigma-Aldrich) as described previously.[9, 35] Equivalent MTT assays were performed on cells cultured in the same ratios of PBS to media, but in the absence of peptide assemblies to confirm that the culture conditions were non-toxic. Absorbance readings of untreated control wells in 100% cell culture media were designated as 100% cell viability. Statistical analysis was performed by one-way ANOVA tests with Tukey comparison in the software GraphPad (Prism) ***p < 0.001.

### 2.10 Amyloid Uptake Experiments

SH-SY5Y cells seeded onto Lab-Tek II imaging dishes (Nunc) 5 × 10^4^ cells per mL and incubated for 24 h. After this time the cells were co-incubated (37 °C, 5 % CO_2_) with peptide assemblies (160 μg mL^-1^) for 48 h. Amytracker 630 (Ebbabiotech) was diluted (1:500) in cell culture media and incubated with the cells for 30 mins. After incubation the cells were washed in cell culture media and imaged using a Ziess LSM 900 confocal microscope using an excitation wavelength of 488 nm, emission was measured at a wavelength of 550 nm, using a x40 objective.

### 2.11 Mitochondrial Respiration Assays

SH-SY5Y were harvested and 5 × 10^4^ cells were seeded into a Seahorse cell culture plate and incubated overnight at 37°C in 5 % CO_2_, to allow them to attach to the surface. The following day, the cells were treated with ILLIIM, RNYIAQVD or AAAAAA assemblies at various concentrations. Untreated cells, as well as cells cultured with equivalent ratios of PBS to media, no peptides, were included as controls. Two days later, the treatment media was removed and 180 µL of XF assay medium pH 7.4 (unbuffered DMEM supplemented with 10 mM glucose, 2 mM sodium pyruvate and 2 mM glutamine) was added.

Mitochondrial respiratory function in live SH-SY5Y cells was measured using the Seahorse XFe96 Extracellular Flux Analyzer (Agilent Technologies, California, USA) via changes in the Oxygen Consumption Rate (OCR) following the sequential addition of pharmacological agents (2 µM oligomycin, 1 µM CCCP, 5µM rotenone, 1µM antimycin A plus 20 µM Hoescst 33342). Between each compound treatment, the average of three measurement cycles of OCR was taken, each cycle including a 3-min mix step and a measurement time of 3 min. Each condition tested had a minimum of three replicate wells, with the average of the values taken per experiment. The final injection of the assay used antimycin A and Hoechst 33342, a cell-permeable DNA dye, that together with a BioTek Cytation 1 imager (Agilent Technologies) allowed for imaging and counting of nuclei per well directly, data used to normalize the Seahorse parameters per cell number.

## 3. Results & Discussion

### 3.1 Solutions of ILLIIM and RNYIAQVD in HFIP rapidly form β*-*sheet rich assemblies when diluted into PBS

In the previous study investigating the self-assembly of ILLIIM (from ORF6) and RNYIAQVD (from ORF10)[9] we used a common protocol for ensuring the monomerization of short hydrophobic peptides that involved heating the peptide solutions, repeated vortexing to dissolve the peptides, followed by a slow cooling to initiate self-assembly.[36–38] Whilst this method did dissolve the majority of peptide, small insoluble seeds remained which have been shown to amplify the formation of new amyloids.[26–28] To eliminate the presence of these insoluble seeds we used the fluorinated alcohol HFIP as a solvent. HFIP is an excellent disruptor of h-bonds and hydrophobic interactions and is frequently used to solubilise amyloidogenic proteins/peptides.[11, 31, 39] In figure 2 we confirm by multiple techniques that when solutions of ILLIIM and RNYIAQVD dissolved in HFIP are diluted into aqueous solutions of PBS they rapidly form assemblies composed predominantly of β*-*sheets. Quantification of the secondary structure contributions (Figure 2a) of the assemblies by non-linear curve fitting of the amide I band of synchrotron FT-IR spectra (Supporting Information, Figure S1) revealed that ILLIIM assemblies are composed of 95 % β*-*assemblies (63 % β-sheet, 16 % β-turn and 15 % from an amyloid specific extended β-sheet network[40]), whilst RNYIAQVD is composed of 82 % β-assemblies (42 % β-sheet, 9 % β-turn and 31 % extended β-sheet network). As expected, the non-amyloid forming control peptide (AAAAAA) is predicted by FT-IR to possess only a small percentage of β-sheet structure indicating that it is likely non-amyloidogenic. Circular Dichroism (CD) spectra of the HFIP assemblies look almost identical to the spectra in Charnley *et. al.*[9] with ILLIIM possessing a classic β-sheet structure with a small minima around 225 nm and a maxima around 200 nm, secondary structure analysis by the BeStSel algorithm [41, 42] predicts that this spectra is composed of a mixture of right and left twisted β-sheets and β-turns (table 1). The CD spectra of RNYIAQVD looks more complex and is likely made-up of multiple contributions. BeStSel predicts this to be a mixture of right-handed β-sheets and α-helices, however the RMSD values for RNYIAQVD are more than 3 times that of ILLIIM suggesting an imperfect fitting. The control peptide consisting of 6 alanine residues is unlikely to have significant ordered secondary structure, and this is reflected by the large RMSD (16.03) when trying to fit the CD spectra in figure 2b, thus any secondary structure predictions from CD spectra of AAAAAA are considered untrustworthy.

**Figure 2:**
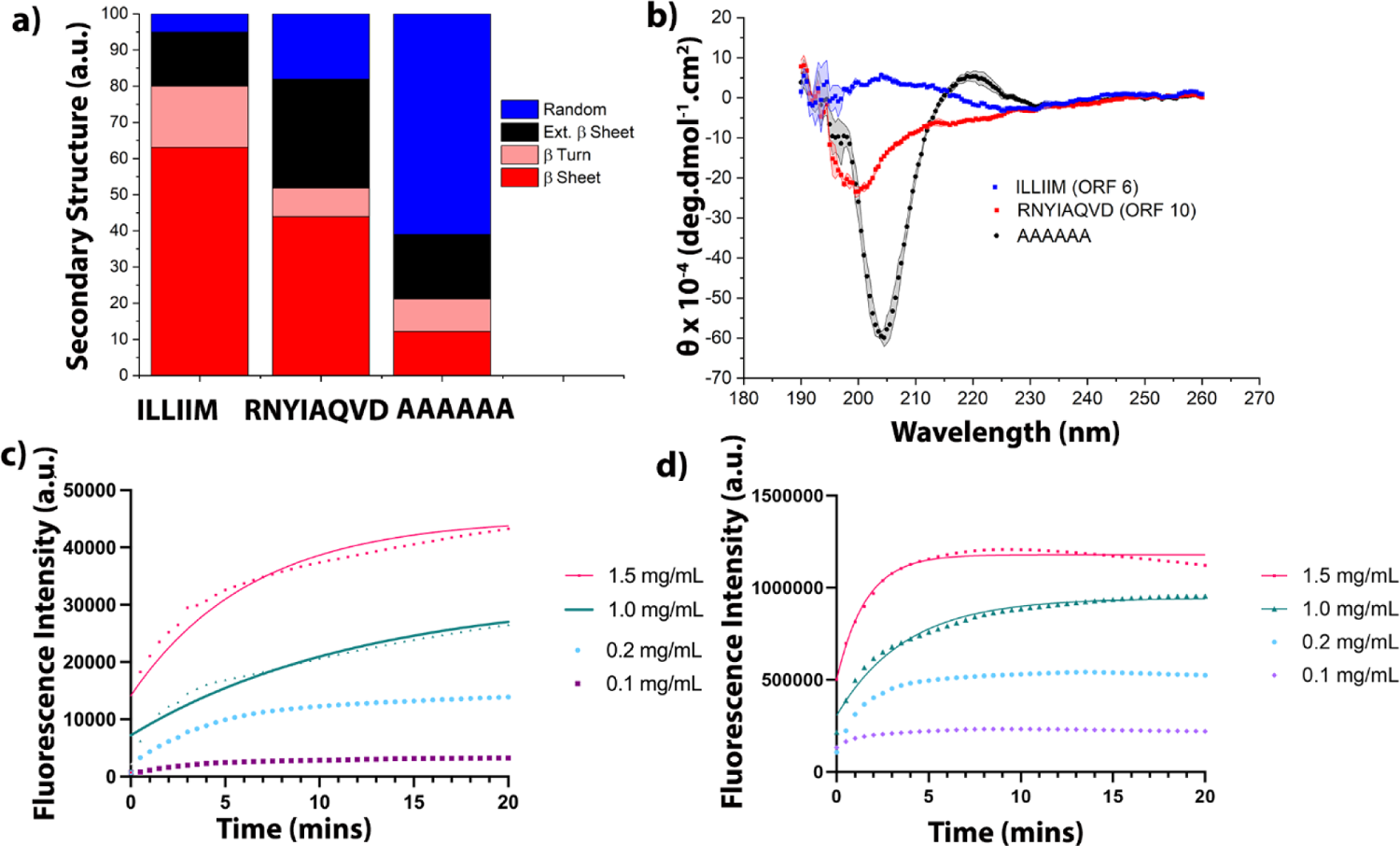
Increased solubilisation in HFIP does not prevent the formation of β-sheet rich assemblies. ILLIIM and RNYIAQVD formed from HFIP solubilised monomers, spontaneously form β-sheet rich assemblies as shown by a) FT-IR, b) CD, c) ThT (ILLIIM), d) ThT (RNYIAQVD).

**Table 1:**
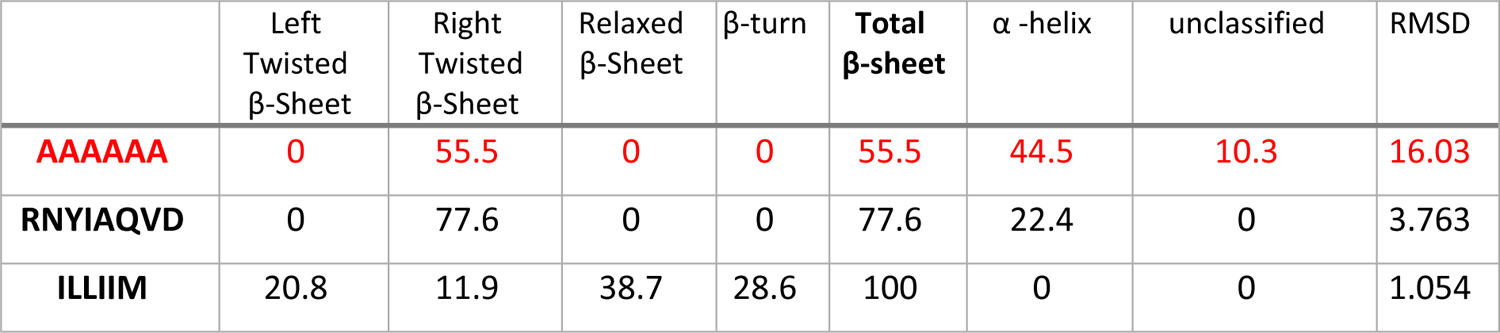
Secondary Structure Determination from CD spectra (results from AAAAAA highlighted in red due to a high error in the RMSD, indicating the results are unreliable)

The kinetics of aggregation of ILLIIM and RNYIAQVD were measured over multiple concentrations for both peptides by Thioflavin T (ThT). Both peptides underwent rapid self-assembly with no observable lag phase as is typical for short amyloidogenic peptides[25, 30, 43]. The kinetics of assembly for both peptides appears to be concentration dependent with the highest concentrations used reaching a plateau in less than 5 mins [1.5 mgmL^-1^]. Curve fitting allowed us to calculate rate constants for the two peptides at the higher concentrations tested. As expected the rate constants increase with increasing concentration, specifically 1.5 mgmL^-1^ ILLIIM assemblies had a rate constant more than 3 times that of 1 mgmL^-1^ assemblies (Table 2).

**Table 2:** Measured Biophysical Properties of the Amyloid Assemblies Studied.

|  | ILLIIM | RNYIAQVD | AAAAAA |
| --- | --- | --- | --- |
| MSED Persistence Length ($\mu\text{m}$) | $13.83 \pm 0.23$ $\mu\text{m}$ | $1.03 \pm 0.047$ $\mu\text{m}$ | no fibers formed |
| Mature Fibril (mode) | 20 nm | 8 nm | no fibers formed |
| Percentage β-sheet (FT-IR) | 63.2 % | 43.6 % | 12.06 % |
| Percentage β-turn (FT-IR) | 16.7 % | 7.8 % | 8.9 % |
| Total β-content (CD) | 100 % | 77.6 % | 55.5 % (large error) |
| Rate constant (ThT @ 1.5 mgmL<sup>-1</sup>) | 0.159 mins<sup>-1</sup> | 0.6253 mins<sup>-1</sup> | No ThT signal |
| Rate constant (ThT @ 1 mgmL<sup>-1</sup>) | 0.083 mins<sup>-1</sup> | 0.271 mins<sup>-1</sup> | No ThT signal |

### 3.2 Increased solubilisation of peptides reduces self-seeding and destabilises the assemblies formed

The spectroscopic data presented in figure 2 suggests that similarly to when assembled in the presence of insoluble seeds the HFIP solubilised peptides form β-sheet rich assemblies. However, when the assemblies are imaged by AFM and TEM we see that the polymorphic landscape of the assemblies is significantly altered. Previously, in the presence of insoluble seeds ILLIIM formed almost exclusively crystalline structures, whereas RNYIAQVD formed a mixture of crystals and twisted fibrils[9]. Now for ILLIIM we still see the presence of crystals, but in addition we see a large population of fibrillar nanostructures (figure 3 a-e, Supporting information, figure S2). High resolution AFM analysis of the ILLIIM assemblies determined that the crystalline assemblies were composed of individual protofilaments 8 nm in diameter (figure 3c-e, and supporting information figure S3 for an enlarged image more clearly showing the individual protofilaments), similar to crystalline assemblies seen for a range of different amyloidogenic peptides[25, 29]. For RNYIAQVD we observed further differences with the preferred architecture formed consisting of a mixture of twisted fibrils (width ∼ 8 nm, figure 3l) and oligomers (figure 3g, supporting information, figure S4) with an average diameter of around 30 nm. For the AAAAAA control peptides we see no evidence of fibril formation, however AFM (Figure 3j) and TEM imaging (Figure 3k) do reveal the presence of amorphous aggregates. Statistical analysis using the opensource software FiberApp[34] (figure 3i) and AFM cross-sections (figure 3l) revealed that the distribution of assembly widths was much smaller for RNYIAQVD compared to ILLIIM with the modal width occurring at 8 nm (Figure 3i), in good agreement with the protofibril width recorded by high resolution AFM of ILLIIM assemblies. This suggests during the assembly of ILLIIM lateral interactions between fibrils are preferred generating 2D crystalline structures composed of close packed protofilaments[25, 29]. RNYIAQVD forms similar sized protofilaments but these filaments have a lesser tendency to laterally associate and form larger multi-filamentous structures. This is likely due to the increased hydrophobicity of ILLIIM, all 6 residues being hydrophobic, therefore the close packing of protofilaments, excluding water molecules, would be preferred compared to RNYIAQVD where only Ile, Ala and Val possess hydrophobic side chains.

**Figure 3:**
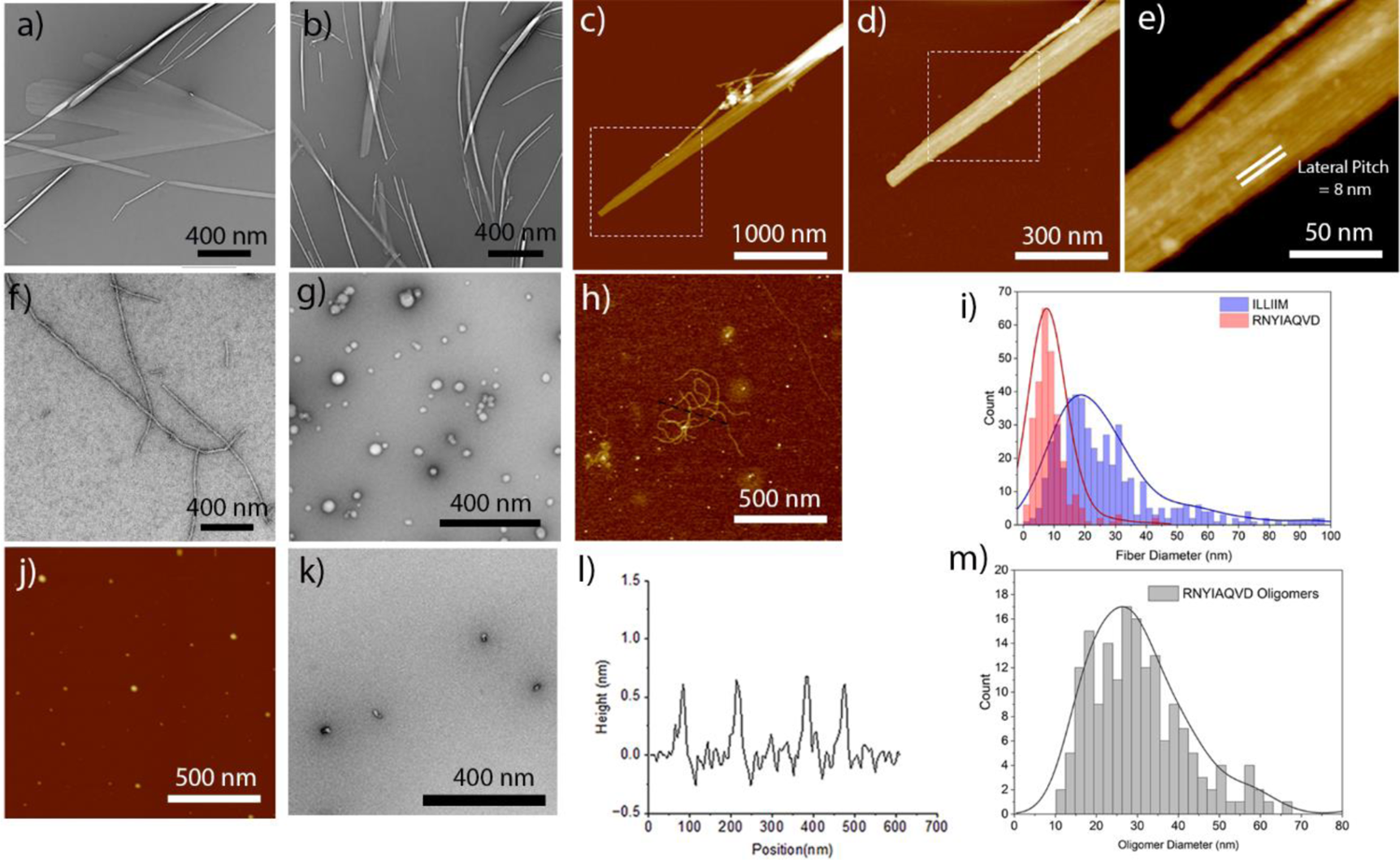
The assemblies formed are highly polymorphic, but overall starting with a highly solubilised starting materials resulted in higher energy assemblies. **a-b)** TEM images of polymorphic crystal and fibrillar assemblies of ILLIIM, c-e) a series of AFM images with increasing magnification showing that ILLIIM crystals are composed of close-packed protofilaments with a diameter of 8 nm, z-scales of (c-e) are 66, 35 and 18 nm respectively (f-g) TEM and (h) AFM images of polymorphic RNYIAQVD assemblies consisting of fibrils (f, h) and oligomeric species (g), AFM (h) has a z-scale of 4 nm, (i) distribution of fibril diameters calculated from a large dataset of TEM images (> 900 fibrils for ILLIIM and RNYIAQVD), j) AFM and k) TEM images of amorphous aggregates formed from the control peptide (AAAAAA), l) line section through the RNYIAQVD fibrils, m) distribution of RNYIAQVD oligomer diameters.

The tendency for ILLIIM to preferentially form less flexible multifilamentous assemblies was confirmed by the persistence length calculations performed in FiberApp where RNYIAQVD was found to have a persistence length of 1.03 ± 0.047 µm (relatively low for semi-flexible amyloid fibrils[25, 30, 44, 45]) whilst ILLIIM has a much larger persistence length of 13.83 ± 0.23 µm due to the increased stiffness of the fibers arising from the strong laterally associated protofibrils (see table 2).

This change in the polymorphic landscape of the assemblies formed in HFIP can be explained by a change in the free energy of the aggregates. It is well-established that amyloids occupy a low energy stable intermediate in the protein folding free energy landscape[32]. More recent research has established that different amyloid polymorphs have differing comparable free energies following the series (oligomers > fibrils > crystals) with amyloid crystals proposed to occupy a global energy minima.[24, 25] It is also well established that insoluble seeds can accelerate/promote the self-assembly of amyloid assemblies[26] thus resulting in lower free energy assemblies, such as crystals and fibrils. The current study supports this, when assembly occurs from better solubilised peptides, we see an increase in the formation of higher free energy assemblies compared to incomplete solubilisation via heating.[9] For ILLIIM we see a reduction in the number of crystals, and a corresponding increase in the number of fibrils, and for RNYIAQVD a reduction in the number of crystals and fibrils and an increase in the number of oligomers (figure 1).

In the cellular space (intra or intercellular) it is likely that amyloid seeds of ORF6 will not be present in high concentrations (excluding the possibility of ORF6 amyloid build-up due to multiple COVID-19 infections), thus the viral amyloids formed may have limited stability. This limited stability may go some way to explain the largely transient nature of the neurological symptoms of COVID-19 compared to NDs such as AD and PD. Further, this finding may offer hope for the possibility of using anti-amyloid therapies similar to monoclonal antibodies like Lecanemab[46] to disassemble the higher energy amyloids formed *in vivo* from SARS-CoV-2 peptides. However, if amyloid seeds of viral protein can accelerate the formation of endogenous amyloids (*e.g.* Aβ, Tau or α-synuclein) *in vivo* this could imply a worrying link between COVID-19 and the accelerated onset of neurodegenerative diseases (NDs) like Alzheimer’s and Parkinson’s. Indeed, others have already demonstrated that viral amyloids can cross-seed and accelerate the formation of endogenous human amyloids.[14] Furthermore, there are indications that severe or repeated infections with SARS-CoV-2 may result in an increase in patients displaying neurodegenerative disease like symptoms,[10, 47–50] and a similar pandemic of Parkinsonism was reported in the early 20^th^ century following the Spanish Flu.[51]

Supporting the nanoscale imaging data in figure 3, X-ray scattering data collected from the SAXS/WAXS beamline at the Australian Synchrotron confirms significant differences in assembly compared to that seen in Charnley et. al.[9] WAXS spectra produced from the HFIP solubilised peptides (figure 4a) show much reduced Bragg reflections, previously the ILLIIM and RNYIAQVD WAXS plots contained multiple Bragg peaks likely due to a high concentration of highly diffracting crystalline polymorphs. In the data from the HFIP solubilised fragments (figure 4a) only two (broadened) Bragg peaks are clearly resolved at 0.56 Å^-1^ and 1.38 Å^-1^ corresponding to the ubiquitous intersheet and intrasheet amyloid reflections at around 11.2 Å and 4.5 Å respectively.[25, 43, 52] The RNYIAQVD and AAAAAA peptides show no resolvable Bragg peaks, likely to be due to the lack of crystalline assemblies present.

**Figure 4:**
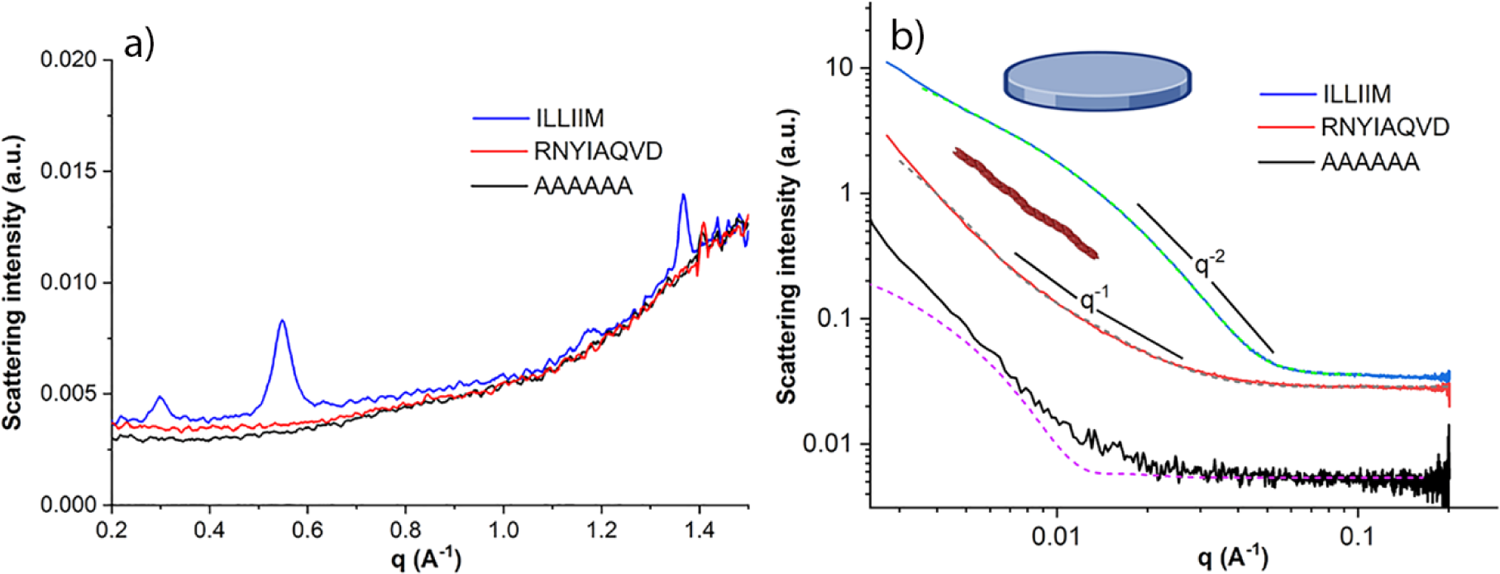
SAXS/WAXS data supports the formation of higher free energy assemblies formed from solubilised monomers. **a)** WAXS spectra of ILLIIM, RNYIAQVD and AAAAAA assemblies, showing characteristic meridional and equatorial Bragg reflections from extended β-sheet networks[52] (typically only detectable crystalline amyloid structures) only for ILLIIM, b) SAXS spectra of ILLIIM, RNYIAQVD and AAAAAA with associated form factor fitting, the SAXS curve of ILLIIM fits to a form factor on a flattened elliptical assembly, the SAXS curve of RNYIAQVD fits to the form factor of an flexible elliptical cylinder, AAAAAA does not fit well to any form factor.

For the ILLIIM SAXS data, as previously, we see a scattering curve with a slope with a q^-2^ dependence consistent with a morphology representing an flattened 2D surface.[9] This is somewhat supported by the fitting of a form factor that describes a flattened sphere (see Supporting information, table S1 for details of form factor fitting), although the best fitting dimensions of this flattened object (38 x 105 nm, 10 nm thick) are not in agreement with the AFM and TEM images in figure 2. This discrepancy suggests that there may be differences in the dominant polymorphs in solution (smaller aggregates) seen by X-ray scattering compared to those most easily imaged by AFM/TEM (larger aggregates may preferentially bind to AFM and TEM imaging substrates). For RNYIAQVD we now see a scattering curve with a q^-1^ scaling in its central region indicative of a high aspect ratio fibrillar structure[30], further this spectra can be well fitted to the form factor of a flexible elliptical cylinder, suggesting that the fibrillar polymorph dominated for RNYIAQVD under these conditions. The AAAAAA polymer scattered very weekly, hence the particularly noisy reduced scattering curve. Whilst no well-fitting form factor could be found for AAAAAA (supporting info, table S1 and figure 4b) the best fit achieved also corresponded to a flexible elliptical cylinder but with a length of less than 150 nm, which is in reasonable agreement with the amorphous aggregates of AAAAAA seen in figures 3 j,k.

### 3.3 Higher energy polymorphs show reduced cytotoxicity

We now turn our attention to the assessment of the biological effects that the SARS-CoV-2 peptide assemblies on SH-5Y5H cells. Thanks to the highly specific binding and low background of the luminescent conjugated oligothiophene (LCO)[53, 54] fluorophore (AmyTracker 630) we were able to observe cell uptake of the amyloid assemblies into cells. After 48 h incubation of the peptide assemblies at 160 µg mL^-1^ we could see clear evidence of Amytracker positive amyloid assemblies (yellow punctate spots) of ILLIIM (Figure 5a) and RNYIAQVD (Figure 5b) in the cytoplasm of the cells. In the case of the control peptide AAAAAA very few punctate spots could be observed suggesting that cell uptake is specific to the amyloid species formed. Analysis of the average number of stained aggregates per cell revealed a similar number of aggregates for ILLIIM and RNYIAQVD (2 per cell) and 0.1 aggregates per cell for the AAAAAA aggregates. We believe this is the first direct microscopic evidence of amyloid fibrils from SARS-CoV-2 viral proteins crossing the cell membrane. This is highly relevant for a number of reasons: first, amyloid fibrils are known to have an almost ubiquitous ability to interact with and disrupt cell membranes[55–57] they are also highly efficient transfection agents[58, 59]. Therefore our results suggest that viral amyloid fibrils could be aiding viral infection and/or cellular escape during the viral replication cycle, suggesting a previously unconsidered role of amyloid in SARS-CoV-2 infection.

**Figure 5:**
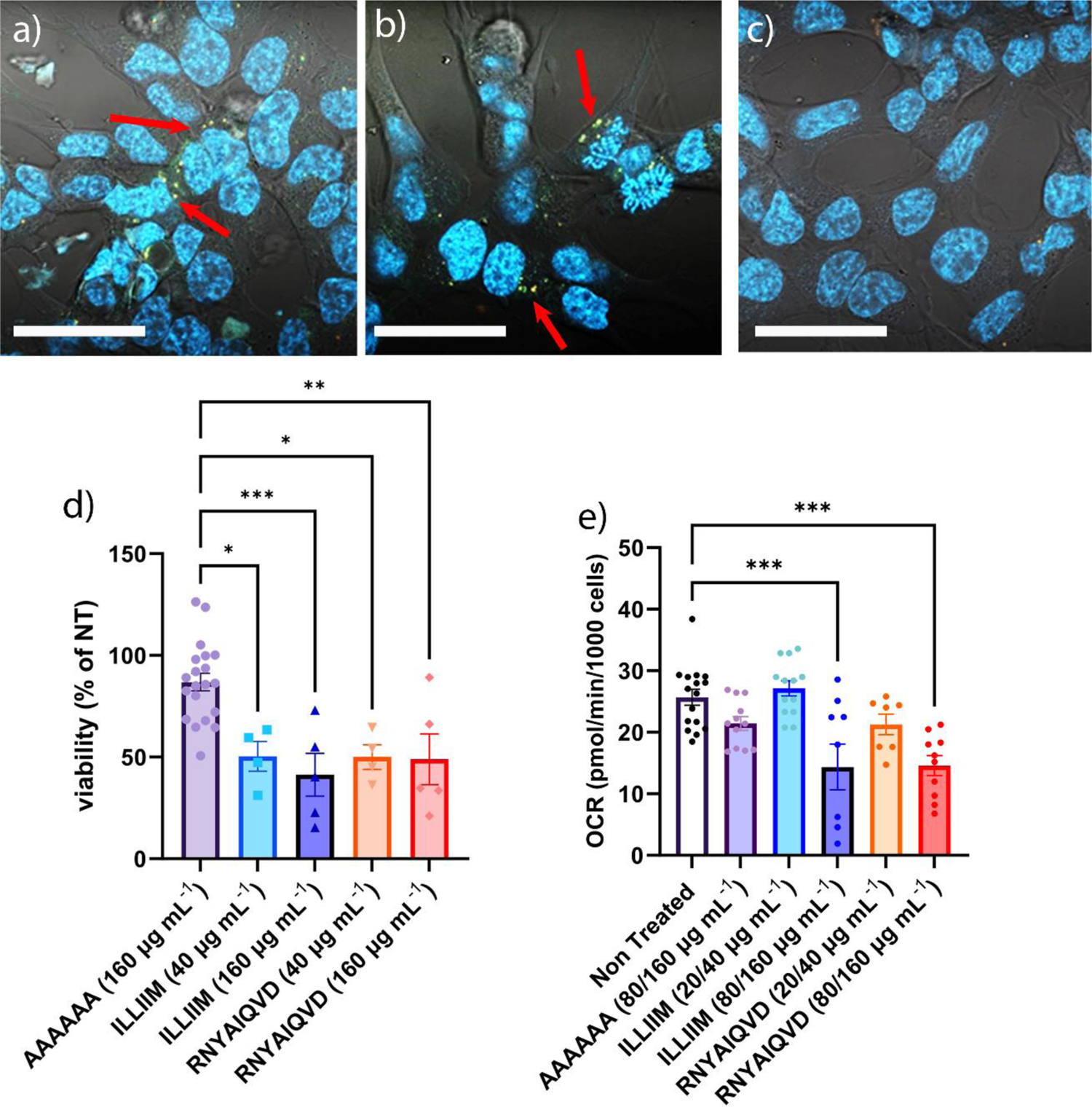
Amyloid aggregates from SARS-CoV-2 peptides readily enter neuronal cells, and this causes a reduction in their viability and their mitochondrial respiration. (a-c) Laser Scanning Confocal Microscopy (LSCM) images of SH-SY5H cells with fluorescently stained amyloid aggregates in the cytoplasm of the cells (a) ILLIIM assemblies, b) RNYIAQVD assemblies, c) AAAAAA assemblies, blue = hoescht nuclear stain, yellow = Amytracker 630 stained amyloid assemblies (as indicated by red arrows), Scale bars = 50 µm. d) MTT cell viability assays for ILLIIM and RNYIAQVD and AAAAAA) relative to a non-treated control, e) basal oxygen consumption rate of the SH-SY5H cells exposed to ILLIIM, RNYIAQVD and AAAAAA. For all graphs the error bars represent the standard error of mean of three of technical replicates recorded over at least four biologically independent replicates. Statistical analysis was performed by one-wat ANOVA with Tukey comparison. *p < 0.1, **p < 0.01 and ***p < 0.001.

In figure 5d we performed MTT assays to measure the reduction in viability of the SH-SY5Y cells after 48 h incubation with the peptide assemblies at a range of concentrations, identified to be relevant in previous studies[9]. Similarly, to Charnley et. al.[9] we observed a statistically significant drop in cell viability at concentrations as low as 40 µg mL^-1^ for both peptides. As expected we see little toxicity for the control AAAAAA peptide which does not form amyloid aggregates (Figures 2-4). In Charnley *et. al.* we speculatively suggested that the increased toxicity seen for ILLIIM compared to RNYIAQVD maybe due to the increased thermodynamic stability of the ILLIIM (crystal) assemblies versus the RNYIAQVD (fibrils) making it harder for the cells to break down the toxic amyloid assemblies.[9] The data shown here further supports this hypothesis, as previously discussed, the increased solubilisation offered by HFIP results in the preferential formation of higher energy/less stable assemblies and this results in a corresponding drop in the levels of toxicity of the assemblies. In Charnley et. al.[9] the amyloid assemblies formed in the presence of insoluble seeds resulted in a 63 % (ILLIIM) and 59 % (RNYIAQVD) reduction in cell viability at 160 µgmL^-1^. In figure 5 the equivalent drop in cell viability is 52 % for both ILLIIM and RNYIAQVD, with a similar drop seen at other concentrations tested. Combined these results suggest that increasing the solubility of the starting materials reduces the concentration of insoluble seeds, reducing the thermodynamic stability of the amyloid assemblies formed. This reduction in stability translates to a reduction in cytotoxicity. This is likely due to cells being able to process less stable amyloid assemblies more easily.

To investigate the cellular responses triggered by the amyloid assemblies in more detail, we performed a series of experiments to quantify changes in mitochondrial respiration. In figure 5e we show changes to the basal oxygen consumption rate (OCR) in the presence of the viral amyloids. To better untangle subtle changes in basal OCR we have collated the data from peptide concentrations at 20 and 40 µg mL^-1^ into a combined low concentration bin and the data at 80 and 160 µg mL^-1^ into a high concentration bin. At high concentrations we see a significant drop in basal OCR for both ILLIIM and RNYIAQVD however, at low concentrations this drop in OCR is absent. This result was somewhat surprising as it does not agree with the data from the MTT assays (Fig 5 d) where significant drops in metabolic activity were observed at all peptide concentrations tested. The effects of amyloid assemblies on mitochondrial respiration are complex and previous investigations have found evidence of both increased[23] and decreased OCR.[20] A speculative explanation as to the lack of significant change in OCR at low (but still cytotoxic, figure 5d) concentrations is that we are seeing the results of two populations of cells within the same sample. One population is undergoing hyperactive mitochondrial respiration as in Ulgade *et. al.*[23] (and therefore has an increased OCR) and a second population consists of dead or dying cells (with much reduced OCRs[20, 21]). At low concentrations the changes to the overall OCR are almost completely cancelled out by the two populations but at higher peptide concentrations when the population of dead or dying cells becomes larger, we see a significant drop in basal OCR (albeit still partially masked by a small population of hyperactive cells). Additional experiments are currently ongoing to untangle the complex relationship between a variety of different amyloid peptides/proteins and their effect on mitochondrial respiration and will be the focus of subsequent works.

To summarise the data presented here, we have shown that by improving the solubility of the peptides we can reduce the presence of insoluble seeds allowing us to study the self-assembly of amyloidogenic viral peptides in more detail. Eliminating the amyloid seeds results in the formation of higher energy polymorphs which, whilst able to cross the cell membrane have less cytotoxic effects on cultured neurons, however we still do observe significant reductions in overall metabolic activity and mitochondrial respiration.

### 3.4 Conclusion

Here we further investigated the ability of fragments of SARS-CoV-2 viral proteins to form amyloid assemblies. We show, for the first time, direct evidence of cellular uptake of viral amyloids into neuronal cells, further supporting the hypothesis that some of the neurological symptoms of COVID-19 and PASC may possess a neurotoxic amyloid aetiology. We also observed that in the absence of insoluble amyloid aggregates, known as seeds, in the starting materials these assemblies form less-stable structures which display reduced toxicity, likely due to them being more easily processed and broken down within the cell. Whilst this data is confined to *in vitro* studies it may offer promise that the neurological effects of COVID-19 and PASC are transient, due to the apparent relative instability of viral amyloids compared to amyloids from neurodegenerative diseases like Alzheimer’s. Further, anti-amyloid therapies may show promise as effective treatments for neuro-COVID and PASC.

## Supporting information

Supplemental Information for Sanislav et. al.

## Acknowledgements

N.P.R. would like to acknowledge The La Trobe Institute of Molecular Sciences (LIMS) for the receipt of a Nicholas Hoogenraad fellowship, and the CASS foundation for partially funding this work through a philanthropic grant (#10053, ‘Determining the role of protein aggregation in COVID-19’). N.P.R would also like to thank Prof. Karla Helbig (LTU) for valuable discussions on the cell types infected by SARS-CoV-2. We would also like to acknowledge that Fig. 1 was created using Biorender.com. This research was undertaken, in part, on the IRM and SAXS/WAXS beamlines at the Australian Synchrotron, part of ANSTO.

