## Supplemental Information for Sanislav et. al. for "Cell invasive amyloid assemblies from SARS-CoV-2 peptides can form multiple polymorphs with varying neurotoxicity"

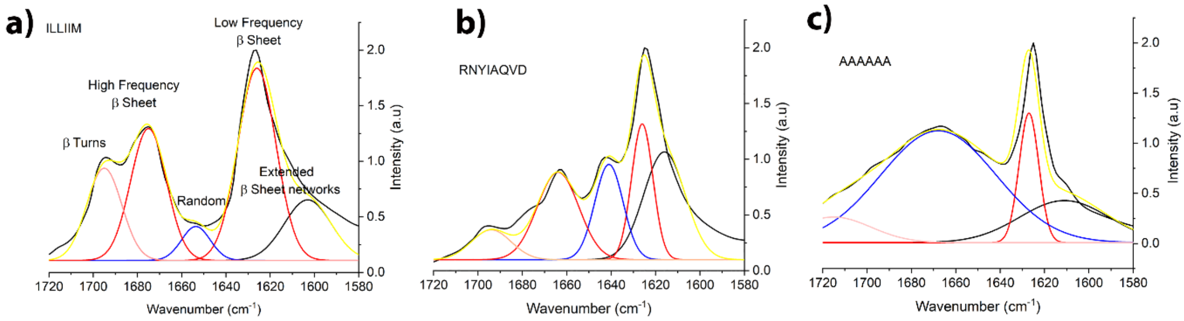


Figure S1: Non-linear curve fitting of amide one band from synchrotron FT-IR spectra of a) ILLIIM, b) RNYIAQVD, c) AAAAAA assemblies. All fitting performed using the non-linear curve fitting tool in Origin Pro 21.

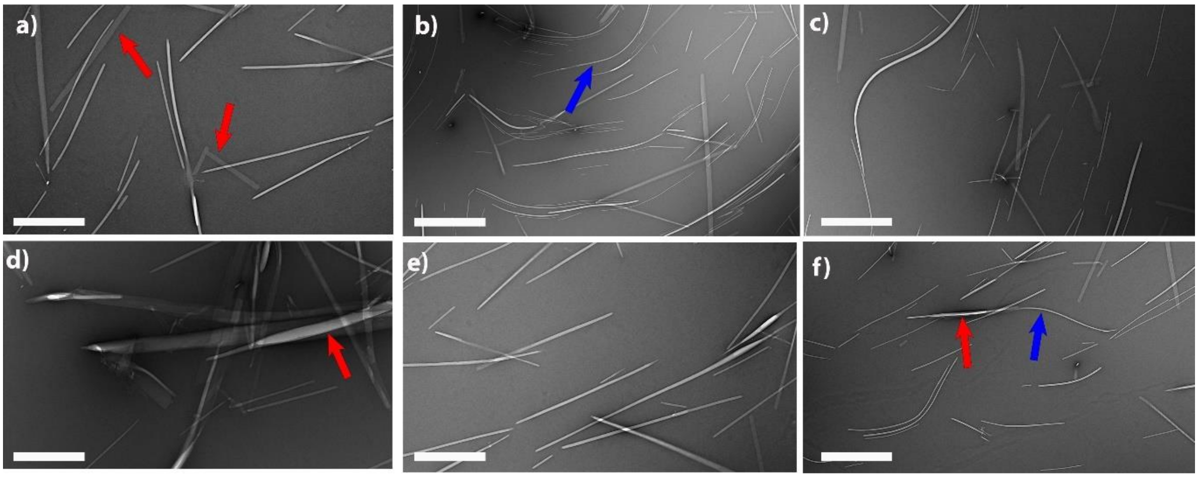


Figure S2: Additional TEM figures of ILLIIM assemblies, red arrows highlight crystalline polymorphs, blue arrows highlight crystalline polymorphs. Scale bars a) = 200 nm, b) = 600 nm, c) = 400 nm, d) = 200 nm, e) = 200 nm, f) = 400 nm

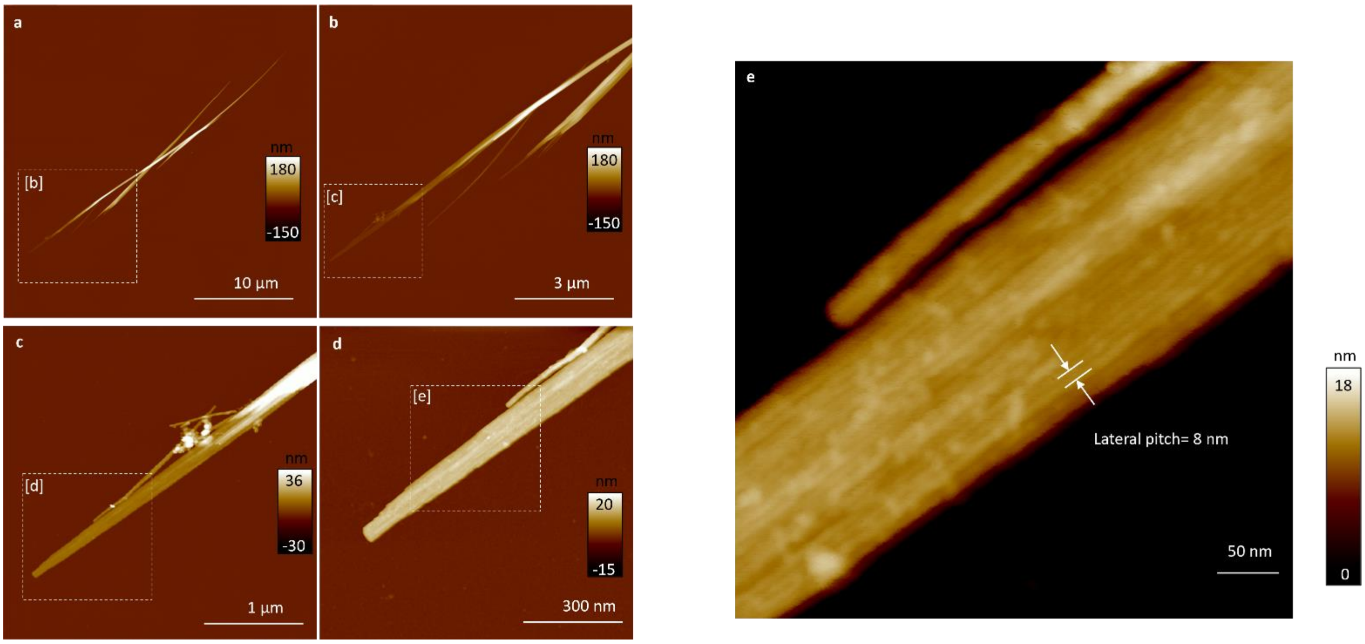


Figure S3: Additional AFM images showing that ILLIIM crystals are made from laterally associated protofilaments with a diameter of 8 nm.

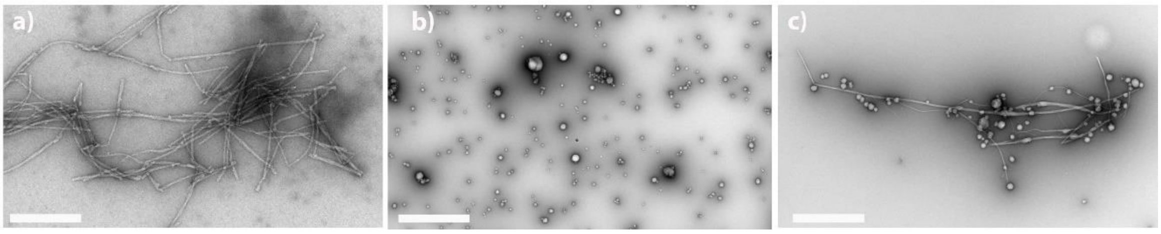


Figure S4: Additional TEM figures of RNYIAQVD assemblies, showing a) fibrillar polymorphs, b) oligomers, c) mixed populations of oligomers and fibrils. Scale bars a) = 200 nm, b) = 1000 nm, c) = 600 nm.

Table S1: Fitting Parameters used to fit SAXS plots in figure 4 (NB. for $\chi^2_R$ a perfect fit = 1)

| | Form Factor Used | Length (nm) | Width (nm) | Thickness (nm) | Reduced Chi2 ($\chi^2_R$) |
|---|---|---|---|---|---|
| ILLIIM | Flattened bicelle | 105.34 | 38.84 | 10 | 0.81 |
| RNYIAQVD | Flexible cylinder | 3.17 x$10^{32}$ (∞) | 50.2 | n/a | 1.15 |
| AAAAAA | Flexible cylinder | 149.6 | 21.9 | n/a | 0.35 bad fit |
